# Self-feeding living materials enabled by cell responsive glycogen nanoparticles as metabolic batteries

**DOI:** 10.64898/2026.06.11.731644

**Authors:** Melvin Gurian, Niels G.A. Willemen, Isa R. Porsul, Nicole Bassous, Jarno Hiemstra, Debby Gawlitta, Su Ryon Shin, Jeroen Leijten

## Abstract

Scaling engineered living materials to clinically relevant dimensions is limited by diffusion-dependent depletion of oxygen and nutrients, which rapidly induces metabolic failure. We introduce glycogen as nutritional nanoparticle that provides cell-mediated, autonomous nutrient release to support long-term survival under extreme metabolic stress. We demonstrate that human mesenchymal stromal cells (hMSCs) survive for weeks in anoxia and serum deprivation when provided extracellular glycogen. Contrary to long-held assumptions, hMSCs secrete glycogen-degrading enzymes, enabling cell-density controlled extracellular glycogenolysis and sustained release of glucose and metabolic intermediates, positioning glycogen as the first-of-its-kind metabolic battery. This cell-responsive process maintains metabolic activity, limits glycolytic acidosis, and enhances pro-angiogenic signaling. To translate this mechanism into a versatile materials platform, we engineered core-shell dextran-tyramine microcapsules that stably encapsulate glycogen while permitting diffusion of enzymes and degradation products. Integrated into centimeter-scale GelMA constructs, these microcapsules maintained hMSC viability and function for at least one month under anoxia. In vivo, glycogen-loaded implants promote deep cellular infiltration, enhanced matrix remodeling, increased M2 macrophage polarization, and orchestrated accelerated vascularization. This work establishes the novel concept of glycogen-based nutritional nanoparticles as metabolic batteries to endow engineered tissues with autonomous self-feeding capacity, enabling scalable and functional living materials for regenerative medicine and related technologies.

## 1. Introduction

The engineering of living materials has numerous high value applications including cultured meat, living soft robotics, drug screening models, and bioartificial organs.^[1–5]^ Although substantial progress has been made in producing viable microscale constructs, scaling living systems beyond the cubic-millimeter range remains a longstanding and unresolved challenge. Specifically larger constructs exacerbate diffusion limitations, leading to oxygen and nutrient deprivation in deeper regions.^[6–8]^ Although hypoxia can stimulate regeneration, anoxia induces cellular dysfunction and necrotic core formation leading to inevitable implant failure.^[1, 3, 6–9]^ Overcoming this challenge represents a key stepping stone towards the engineering of clinically and industrially relevant living materials capable of maintaining long-term viability and function.

Historically, the main driver of necrosis in engineered large living matter was assumed to be lack of oxygen, which led to the development of various oxygen releasing materials.^[3, 10]^ The most commonly explored strategy to generate oxygen in situ is based on encapsulating solid peroxides in hydrophobic (micro)materials, thereby producing hydrogen peroxide as an intermediate product.^[10]^ Despite the ability of these materials to maintain tissue viability for two weeks, and even promote angiogenesis in vivo, their clinical translation is hindered by cytotoxic hydrogen peroxide release as well as their dependency on (xenogeneic) enzymes or metalorganic frameworks to reduce the resulting cytotoxicity.^[11, 12]^

It has recently been hypothesized that lack of oxygen is not directly lethal but instead initiates rapid nutrient depletion by forcing cells to rely on anaerobic respiration, which ultimately leads to starvation-induced cell death. Providing oxygen deprived cells with constant access to nutrients in culture media has been reported to prevent ‘anoxia-induced’ cell death.^[13–17]^ Yet, development of nutrient-releasing materials has remained surprisingly underexplored. Specifically, strategies to achieve nutrient release have remained rare and currently rely on short-lived passive glucose release from carrier materials ^[18]^ or release from plant-based polysaccharides that either rely on outside-in diffusion of systemic enzymes such as amylase or on supplementation of xenogeneic enzymes with short half-lives, which associate with spatially restricted and/or short-lived glycose release.^[19, 20]^ The transient reliance and dependence on non-human enzymes limits the function and translational potential of the few approaches that currently exists, highlighting the need for a fundamentally new strategy to engineer long-term self-feeding living materials.

Here, we introduce the novel strategy of using glycogen as a nutritional nanoparticle for the engineering of autonomous long-term self-feeding living materials. Glycogen is a natural dendrimer nanoparticle, composed of annealed glucose molecules around a single nucleated glycogenin molecule, which has naturally evolved as an intracellular energy reservoir for the reversible storing of glucose via enzymatic reactions.^[21–23]^ As glycogen is an essential part of human physiology, it has been explored for various biomedical applications. For example, it has been used as a drug delivery vehicle, vaccine adjuvant, and contrast agent,^[24]^ which has confirmed its high biocompatibility, ease of functionalization ^[24, 25]^ and ability for safe renal clearance.^[26]^ However, because glycogen degradation is historically assumed to occur only intracellularly, glycogen’s ability to act as an extracellular material system to release glucose has remained unexplored. Surprisingly, we here demonstrate that human cells secrete the enzymes required to extracellularly digest exogenous glycogen nanoparticles, enabling cell-triggered release of glucose at levels sufficient to maintain their survival under chronic anoxia for clinically relevant durations. Additionally, we introduce glycogen microcapsules as metabolic batteries that provide a platform strategy to endow any living material with self-feeding abilities in a scalable and translatable manner.

## 2. Results and discussion

### 2.1 Glycogen maintains cell functionality under anoxia for clinically relevant time periods

When creating large living materials, diffusion limitations create an anoxic environment that causes massive cell death.^[16]^ Specifically, cells exposed to anoxia drastically enhance their nutrient consumption in order to compensate for their inefficient ATP production, which causes rapid microenvironmental nutrient depletion and subsequent starvation-induced cell death (**Fig 1a**). Despite starvation being a well-known driver of cell death and implant failure, the specific nutrients required to sustain cell viability under oxygen deprivation remain surprisingly underexplored.

**Figure 1.**
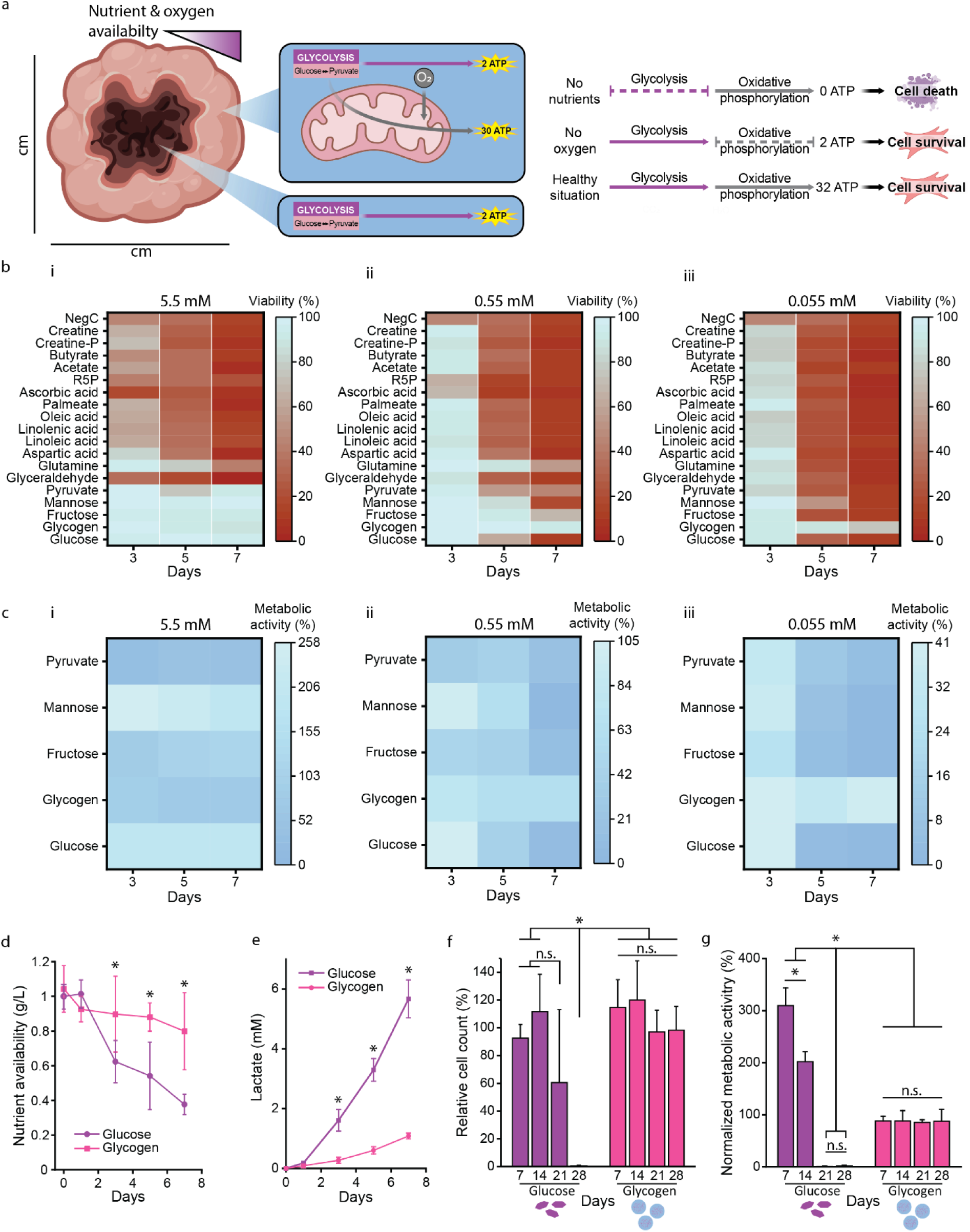
Glycogen maintains cell functionality under anoxia for clinically relevant time periods. a) Schematic representation of cellular respiration that indicates that cell viability is dependent on nutrient availability and that large quantities of nutrients are required in oxygen deprived environments, such as found in large (engineered) tissues. b) Viability and c) metabolic activity of hMSC under anoxia in a nutrient- and serum-free culture medium containing only a single nutrient at different molarities i) 5.5 mᴍ, ii) 0.55 mᴍ, and iii) 0.055 mᴍ. d) Nutrient content and e) lactate buildup in the supernatant of glucose and glycogen supplemented cells cultured under anoxia. f) Relative viable cell counts and g) metabolic activity of hMSCs cultured with 1 g L^-1^ (which equals 5.5 mᴍ) glucose or glycogen in nutrient- and serum-free medium without medium exchange for 28 days in anoxia. n ≥ 3 Significance is indicated with * p < 0.05, and n.s. for p > 0.05, one-way ANOVA with Tukey post hoc test.

To elucidate this knowledge gap, human bone-marrow derived mesenchymal stromal cells (hMSCs) were cultured in chemically-defined nutrient- and serum-free medium under anoxic conditions for up to seven days. We systematically assessed the effects of supplying individual nutrient types on cell survival. Each nutrient type was supplemented at a range of distinct concentrations to an otherwise nutrient- and serum-free medium. To this end, we designed a focused library of nutrients that was organized in different nutrient groups: sugars (glucose,^[27]^ glycogen,^[28]^ mannose,^[29]^ fructose ^[30]^), fatty acids (palmitic acid,^[31]^ linoleic acid, linolenic acid, oleate ^[32, 33]^), amino acids (glutamine,^[34, 35]^ aspartic acid ^[36]^), vitamins (ascorbic acid ^[37]^) and others (creatine,^[38]^ creatine-phosphate, ^[39]^ acetate ^[35]^). Three molarities were tested for each nutrient. For all nutrients except fatty acids and glycogen, the highest molarity was set at 5.5 mᴍ, which corresponds to the normal molarity of glucose in human blood.^[40]^ The highest molarity of fatty acids was set at 300 µᴍ as higher molarities are known to be cytotoxic.^[41]^ For glycogen, concentrations were adjusted to provide an equivalent of 5.5 mᴍ glucose molecules. The two lower molarities were set at a 10-fold and 100-fold lower, representing moderate and severe nutrient depletion. From all nutrient types, only sugars consistently maintained high cell viability (>80%) at the 5.5 mᴍ concentration for all its tested nutrients during the seven days of anoxic culture (**Figure 1b)**. This was notable given that mannose is widely considered as a difficult to metabolize sugar for most human cells, and pyruvate is conventionally assumed to require oxygen to support ATP generation.^[42]^ Most other nutrient classes exhibited higher viability at 0.55 mᴍ than at 5.5 mᴍ after three days, suggesting that elevated concentrations of these nutrients may exert cytotoxic or metabolically stressful effects. This concentration-dependent toxicity limits their feasibility as long-term nutrient reservoirs, as they cannot be safely stored at high concentrations within living materials. Strikingly, among all nutrients tested, only glycogen maintained high hMSC viability throughout the full seven-day anoxic period. The positive effect of glycogen through extracellular supplementation was highly surprising. Specifically, glycogen degradation and its utilization as nutrient requires enzymatic digestion. It is known that blood and serum contain glycogen degrading enzymes such as amylase, which are secreted by specialized cells (i.e. acinar cells) found in the pancreas,^[43]^ and salivary glands.^[44]^ However, due to the serum-free nature of the culture, the glycogen degradation had to be mediated by the hMSCs. As there currently is no current literature that reports that hMSCs are capable of secreting carbohydrate digesting enzymes, we next investigated whether hMSCs could internalize glycogen to allow for intracellular metabolization. Although glycogen has previously been shown to be efficiently internalized by professional phagocytic cells (e.g., macrophages),^[45, 46]^ only a small minority of the hMSC population internalized glycogen, which is likely explained by the non-professional phagocytic nature of hMSCs **(Figure S1a-c)**. In stark contrast, glycogen was able to maintain the viability of the vast majority of hMSCs at molarities as low as 0.055 mᴍ for at least seven days of anoxic culture, strongly suggesting that not internalization but a previously unknown mechanism was behind glycogen’s ability to sustain hMSC metabolism. Regardless, glycogen was able to maintain cell viability at molarities as low as 0.055 mᴍ for at least seven days of anoxic culture.

We hypothesized that glycogen sustains long-term hMSC viability by supporting a lower metabolic rate, enabled by its slow yet continuous release of glucose. Notably, hMSCs supplemented with high levels of simple sugars (e.g., glucose, fructose, and mannose) exhibited markedly elevated metabolic activity compared to control cultures **(Figure 1c i)**. This increase reflects enhanced conversion of resazurin to resorufin, a process dependent on NADH generation, which rises with glycolytic flux, which is amplified under anoxic conditions.^[35, 47, 48]^ Interestingly, glycogen-supplemented hMSC cultures did not show this increase in metabolic activity when supplemented at 5.5 mᴍ, but remained at levels comparable to control cultures. Decreasing the initial nutrient concentration reduced metabolic activity for all sugars relative to both day zero and their higher-concentration counterparts **(Figure 1c ii, iii)**. However, at 0.55 mᴍ or 0.055 mᴍ, only glycogen-supplemented cells maintained measurable levels of metabolic activity throughout the seven day anoxic culture period. This highlights that glycogen can act as a single source of nutrition capable of maintaining cell viability by stably supporting basic metabolic functionality under anoxic conditions, even when present at exceedingly low molarities.

The high potential of glycogen to preserve cell viability even at low concentrations may be explained by its low but consistent conversion rate. While glucose is rapidly internalized and metabolized by cells,^[49]^ glycogen must be enzymatically degraded into glucose prior to its cellular uptake and metabolization.^[50]^ Indeed, while the concentration of supplemented glucose rapidly decreased over time, glycogen only decreased gradually over time **(Figure 1d)**. Consistent with this, lactate buildup occurred at a much lower rate for glycogen supplemented cultures, which is a critical advantage for preventing acidosis within engineered living materials **(Figure 1e)**. Based on these observations, we reasoned that glycogen could support long-term culture without external intervention (e.g., media changes). To this end, hMSCs were cultured in nutrient- and serum-free medium supplemented with either glycogen or glucose, without any medium changes, under anoxic conditions for one month. While metabolic activity rapidly decreased after seven days of culture, glycogen supplemented cultures maintained stable metabolic activity for at least a month **(Figure 1f, g)**. In combination with the serum-free conditions and the concurrent lack of amylase or other supplemented glycogen degrading enzymes, and long-term stability of glycogen under physiological conditions (Figure S1d), these results highlight glycogen nanoparticles as a unique metabolic supplement capable of sustaining cell viability and metabolic activity over clinically relevant timescales, while simultaneously mitigating glycolysis-induced acidosis.

### 2.2 Exogenous glycogen degradation towards glucose is cell mediated and dose dependent

Although it has been textbook knowledge since 1857 that glycogen nanoparticles are the internal glucose storage of cells,^[51, 52]^ glycogen has surprisingly never been explored as an extracellular supplement to maintain cell viability. Typically, glycogen is degraded intracellularly into glucose in a stepwise series of enzymatic reactions, with glucose-1-phosphate (G1P) and glucose-6-phosphate (G6P) being the intermediates ^[50]^ **(Figure 2a)**. To investigate whether the degradation of exogenous extracellular glycogen was initiated through the same process, glycogen phosphorylase (GP), the rate-limiting enzyme for cytosolic glycogen degradation ^[53]^, was blocked using KB-228 in cultures containing different glycogen concentrations. This revealed that inhibiting GP resulted in a significant drop in cellular viability **(Figure 2b, c)**. Moreover, analysis of the supernatant revealed that GP was secreted by hMSCs in a glycogen dose-dependent manner, indicating that the cells can sense the available levels of extracellular glycogen and release GP accordingly **(Figure 2d)**. Furthermore, the concentration of secreted GP is dependent on the cell density **(Figure 2e)** as well as the duration of exposure to glycogen **(Figure 2f)**. Notably, most GP was secreted within the first day of culture, which might be a stress reaction of the cells to the harsh anoxic environment, which is in line with previous research reporting that large quantities of GP are secreted following a stroke or heart attack. Thus, the release of glycogen degrading enzymes may be a natural cellular protection mechanism upon nutrient or oxygen depletion.^[54, 55]^ Additionally, the secretion of GP is not limited to hMSCs, but is present in a wide variety of cell types that are commonly used in tissue engineering and regenerative medicine **(Figure S2a, b)**, indicating the wide applicability of this system. Furthermore, the secreted GP was active and led to the production of G1P and G6P when the supernatant was added to glycogen **(Figure 2g)**. These degradation products also scaled with the initial glycogen concentration during cell culture, which further highlights the concentration dependent secretion of glycogen phosphorylation by the cells. These degradation products (e.g., metabolites) were likewise able to maintain cell viability and metabolic activity under anoxic conditions as demonstrated via exogenous supplementation **(Figure 2h-j)**. Collectively, these findings demonstrated that cells can enzymatically degrade extracellularly glycogen into glucose, enabling sustained cell survival, metabolic activity, and GP secretion. Consequently, this establishes glycogen as the first strategy to provide on demand, autonomous and long-term nutrient delivery in living materials **(Figure 2k)**.

**Figure 2.**
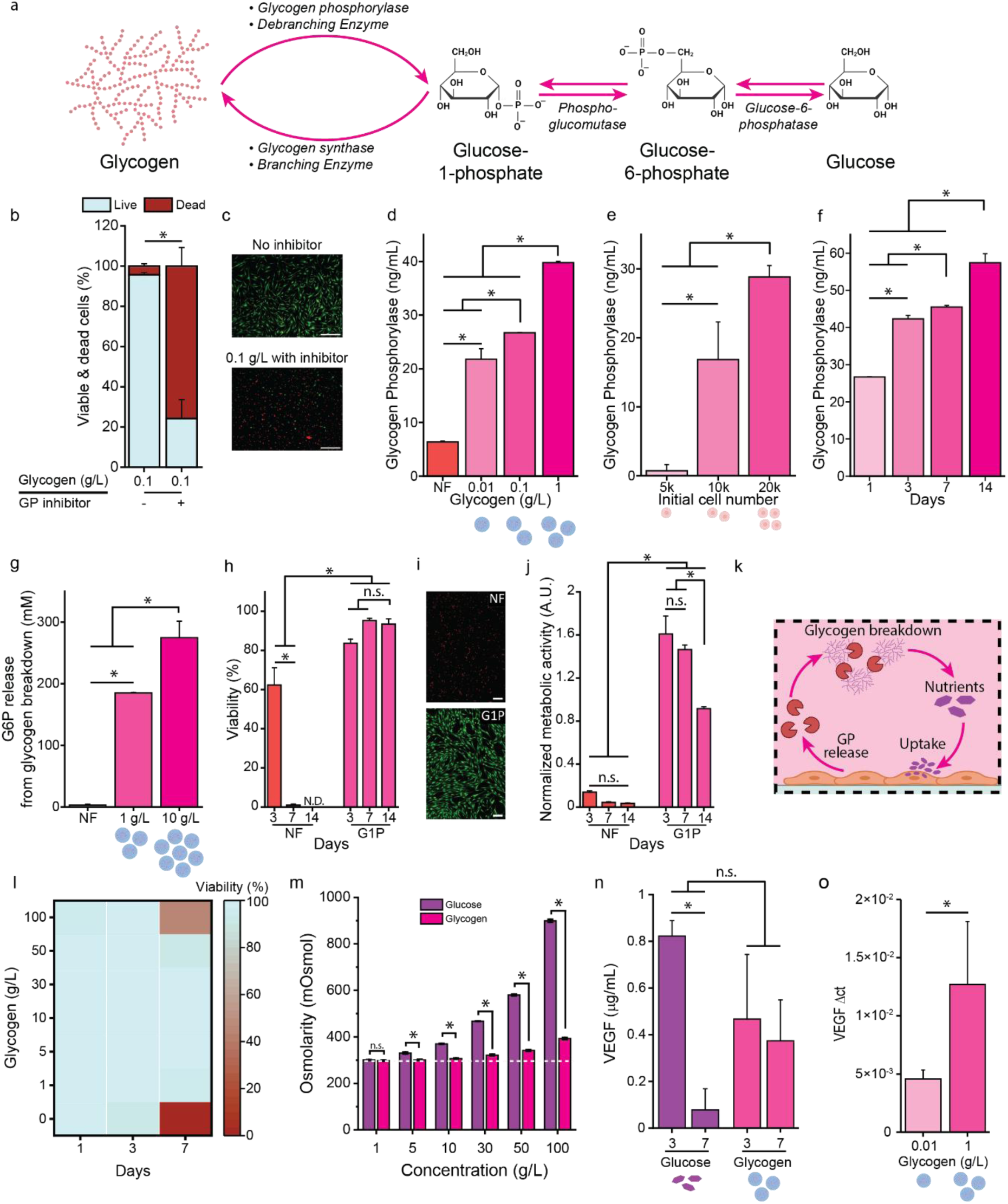
Exogenous glycogen degradation towards glucose is cell-mediated and dose-dependent. a) Schematic representation of endogenous glycogen degradation ^[58]^. b) hMSC viability after 72h while glycogen phosphorylase (GP) activity is being blocked with c) representative live dead images. Scale bars: 200 µm d) GP release of hMSCs exposed to various glycogen concentrations after 24 hours. e) GP release at different hMSC seeding densities after 24 hours. f) GP release over time of hMSCs initially exposed to 0.1 g L^-1^ glycogen. g) Glucose-6 phosphate (G6P) release after GP addition to various glycogen concentrations. NF: Nutrient free medium h) hMSC viability, i) representative images. scale bars: 100 µm, and j) metabolic activity after glucose-1-phosphate (G1P) supplementation. k) Schematic representation of the self-feeding mechanism achieved with exogenous glycogen supplementation. l) hMSC viability after exposure to various glycogen concentrations. m) Osmolarity of PBS supplemented with varying glucose and glycogen concentrations. n) VEGF secretion of cells supplemented with 0.1 g L^-1^ glucose and glycogen under anoxia. o) VEGF expression of hMSC with different glycogen concentrations. n = 3. Significance is indicated with * p < 0.05, and n.s. for p > 0.05, one-way ANOVA with Tukey post hoc test when normally distributed samples, otherwise Kruskal-Wallis.

To investigate the limitations of glycogen supplementation, hMSCs were cultured under anoxia in nutrient- and serum-free medium supplemented with various glycogen concentrations ranging up to 100 g L^-1^ for seven days. Interestingly, hMSC viability remained high for concentrations up to 50 g L^-1^ **(Figure 2l)**. This is in stark contrast to simple sugars such as glucose, which cause severe cytotoxicity at significantly lower concentrations **(Figure S2c)**. This difference was attributed to the low osmotic load of glycogen as compared to other sugars such as glucose **(Figure 2m, Figure S2d-f).** Advantageously, this permits high density storage of glycogen, enabling the integration of vast energy stores into engineered living materials without perturbing local osmolality or inducing cellular (dys)function.

Although we demonstrated that glycogen could maintain cell viability and metabolic activity, it remained unknown if glycogen could support more advanced cellular functions such as cellular orchestration of its microenvironment. As vascularization of nutrient deprived living matter is of essential importance for long-term survival of living implants, we investigated the ability of glycogen supplemented hMSCs to secrete pro-angiogenic factors. Cells naturally produce and secrete pro-angiogenic factors, such as VEGF, under hypoxic or anoxic conditions to facilitate vascularization ^[56]^. To this end, the VEGF secretion of hMSCs cultured in chemically-defined, nutrient- and serum-free medium supplemented with either glucose or glycogen was quantified for up to seven days of culture. While VEGF secretion was initially highest in glucose supplemented conditions, secretion levels had declined sharply after seven days **(Figure 2n)**. In contrast, glycogen supplementation supported stable and sustained VEGF secretion for the entire duration of the experiment, offering a facile approach to program consistent long-term VEGF production and secretion by cells. Additionally, VEGF expression was concentration-dependent on the glycogen concentration (**Figure 2o**). The high energy demand for these processes in combination with the extended starvation of the cells and the continuous VEGF secretion over time strongly suggests that glycogen offers a continuous nutrient source to maintain cell functionality.^[57]^ This was further confirmed by the preserved ability of the hMSCs to differentiate towards adipogenic and osteogenic lineage after 28 days of anoxic culture in glycogen supplemented nutrient- and serum-free medium **(Figure S2g, h)**. Ultimately, these results highlight that glycogen not only maintains basic viability and metabolic activity under anoxia but more advanced cellular functions, such as pro-angiogenic behavior, highlighting the potential of metabolic materials in tissue engineering and regenerative medicine.

### 2.3 Glycogen loaded microgels as metabolic batteries for engineered living materials

To maintain long-term glucose release from glycogen inside engineered living materials, it is essential to also ensure its retention within the construct. To achieve this, the hydrodynamic mesh size of the engineered construct must be smaller than the size of glycogen to avoid passive diffusion. Conversely, because enzymatic degradation of glycogen demands space for conformational changes and enzyme access, the construct’s hydrodynamic mesh size must be sufficiently large to facilitate this process. Together, these opposing constraints severely restrict the material design space. To overcome this limitation, we designed a modular construct composed of glycogen-loaded microcapsules within an engineered hydrogel matrix **(Figure 3a)**. Specifically, the microcapsule shell (e.g., hollow microgel) should prevent glycogen diffusion and allow for enzyme diffusion, while its liquid core should provide the physical space for glycogen deformation. Moreover, the modular design allows the microgels to be incorporated into virtually any biomaterial, offering a universal and customizable material platform.

**Figure 3.**
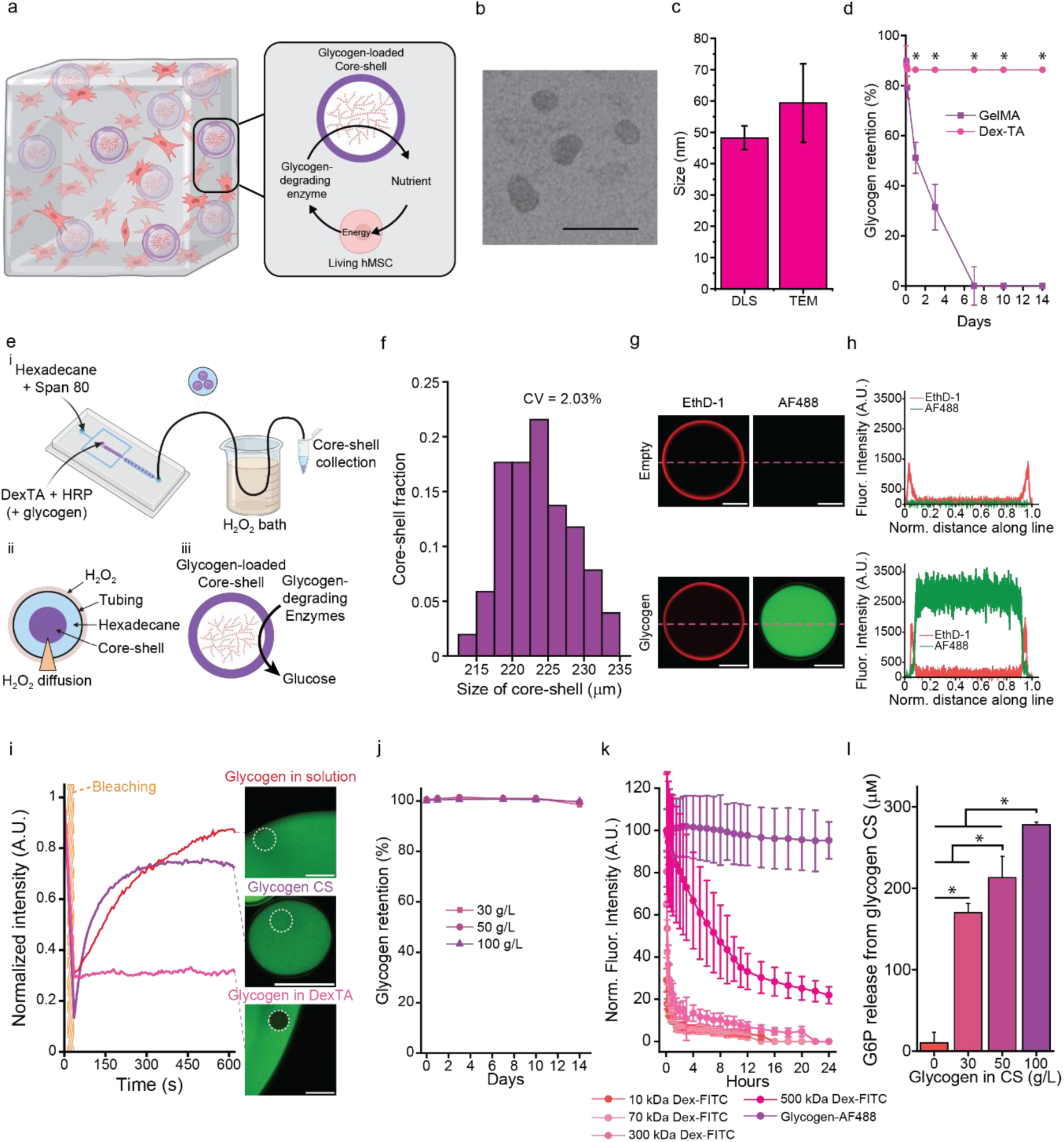
Glycogen loaded Dex-TA microcapsules allow for glycogen retention and allow for long-term enzymatic release of carbohydrate nutrients. a) Schematic representation of a self-feeding large engineered construct based on the incorporation of glycogen loaded microgels. b) Representative transmission electron microscopy (TEM) image of oyster glycogen. Scale bar: 100 nm c) Size of oyster glycogen nanoparticles measured with dynamic light scattering (DLS) and TEM. d) Glycogen retention in hydrogels with large (GelMA) and small (Dex-TA) hydrodynamic radius. e) i) Schematic representation of the Dex-TA core-shell microgel microfluidic production process. ii) The diffusion-based crosslink using H_2_O_2_ enables a timed outside in crosslink that results in iii) core-shell microgels that retain glycogen while allowing its enzymatic degradation and the consequent nutrient release. f) size distribution of glycogen containing core-shell microgels. g) representative images of core-shell microgels where Dex-TA was stained red with ethidium homodimer-1 (EthD-1) and fluorescently labelled glycogen-AF488 was used. scale bars: 100 µm h) with representative intensity profiles. i) Recovery times of ConA stained glycogen in solution, Dex-TA hydrogel, and Dex-TA microcapsules with representative images. Scale bars: 200 µm. j) Retention of various glycogen concentrations in Dex-TA core-shell microgels k) Retention of different sized molecules in comparison to glycogen. l) Glucose-6-phosphate (G6P) release from glycogen loaded core-shell microcapsules after the addition of glycogen phosphorylase (GP) containing cell supernatant. n = 3. Significance is indicated with * p < 0.05, and n.s. for p > 0.05, one-way ANOVA with Tukey post hoc test.

Transmission electron microscopy (TEM) confirmed that glycogen exhibits a nanoparticle-like morphology **(Figure 3b)**, with an average particle size of 59.3 ± 12.5 nm, a finding further supported by dynamic light scattering (DLS) (48.2 ± 3.8 nm) **(Figure 3c)**. As expected, when glycogen was incorporated into a nanoporous hydrogel with a hydrodynamic mesh size smaller than glycogen’s molecular dimension (e.g., tyramine conjugated dextran; Dex-TA) it was retained, whereas it readily diffused out of macroporous hydrogels (e.g., gelatin methacryloyl; GelMA) **(Figure 3d)**. Moreover, although glycogen was entrapped in the Dex-TA hydrogel, no glucose was released, as the hydrogel did not provide the physical space required for enzymatic digestion of glycogen **(Figure S3a)**. Based on this, we hypothesized that encapsulating glycogen within a Dex-TA microcapsule would allow for glycogen retention, even when loaded into macroporous hydrogels such as GelMA, while simultaneously offering the physical space for enzymatic digestion. To this end, glycogen-loaded Dex-TA microcapsules were produced using a microfluidic droplet generator that facilitated sol-gel transition of polymer solution-droplets via diffusion-based outside-in crosslinking ^[59]^ **(Figure 3e)**. This method produced monodisperse populations of glycogen-loaded microcapsules with a diameter of 220.3 ± 3.5 µm **(Figure 3f).** Furthermore, microcapsule diameter could be accurately and predictably tuned by varying the oil-to-aqueous flow rates ratio within the droplet generator **(Figure S3b)**. The addition of glycogen had no notable influence on core-shell microgel production or stiffness, and only slightly increased microcapsule diameter at aqueous flow rates above 10 µL min^-1^. Confocal analysis of ethidium homodimer-1 stained Dex-TA microcapsules revealed a shell thickness of 24 ± 2.1 µm and a non-crosslinked liquid core of 196.3 ± 5.6 µm **(Figure 3g, h, S3c)**. Presence of glycogen within the cores of glycogen-loaded microgels was confirmed using glycogen that was covalently labelled with a fluorophore (**Figure 3g, h).** Additionally, the absence of glycogen within the Dex-TA shell showed that the glycogen and associated proteins ^[60]^ that are often found in oyster glycogen did not interact or interfere with the crosslinking of the hydrogel shell. Fluorescence recovery after photobleaching (FRAP) confirmed glycogen mobility within Dex-TA microcapsules as well as glycogen’s inability to diffuse through a Dex-TA hydrogel **(Figure 3i)**. Indeed, glycogen was stably retained within the microcapsules across all tested concentrations for at least 14 days **(Figure 3j)**. Fluorescence intensity measurements of encapsulated labeled glycogen further showcased a high encapsulation efficiency that increased with increasing glycogen concentration. Specifically, the loss of absolute glycogen for 30 g L^-1^ and 100 g L^-1^ was comparable, further showcasing the predictability and control of and over the system **(Figure S3d)**.

For the entrapped glycogen to support the metabolism of living materials, smaller molecules such as glycogen degrading enzymes as well as glucose must be able to freely diffuse through the microcapsule shell. To evaluate this, FITC-conjugated dextrans of various molecular weights (10 – 500 kDa), representing the size range of relevant small molecules and enzymes, were encapsulated and monitored for their ability to diffusion out over time. This revealed that, unlike glycogen, all tested fluorescent dextrans rapidly diffused through the microcapsule shell **(Figure 3k).** As GP has a molecular weight of ∼200 kDa,^[61]^ it was anticipated to freely diffuse into the microcapsules. The ability of GP to freely diffuse into the microcapsules to degrade the entrapped glycogen was confirmed by incubating the glycogen-containing microcapsules with cell supernatant that contained GP, which subsequently triggered the release of glycogen associated degradation products in G1P and G6P **(Figure 3l)**. Advantageously, G6P release positively correlated with encapsulated glycogen concentration, confirming that release kinetics can be programmed by tuning the concentration of the glycogen stores. Ultimately, these results demonstrate that Dex-TA microcapsules effectively prevent glycogen diffusion while allowing penetration of glycogen-degrading enzymes, thereby functioning as modular metabolic batteries that provide carbohydrate reservoirs for controlled long-term release of nutrients within engineered living materials.

### 2.4 Engineering self-feeding engineered living materials with glycogen as metabolic batteries

To investigate the potential of glycogen-loaded core-shell microgels as metabolic batteries to enable long-term self-feeding of engineered living materials, we evaluated their ability to maintain hMSC functionality under anoxia. Glycogen-loaded or empty microcapsules were added onto transwell inserts that were subsequently introduced on top of hMSC monolayers cultured under anoxia in nutrient- and serum-free medium for 28 days **(Figure 4a)**. Addition of glycogen-containing microcapsules maintained high viability (>80%) for the entire duration of the experiment, while virtually no living cell could be detected while using glycogen-free microcapsules or microcapsule-free controls after only seven days of culture **(Figure 4b)**. Not surprisingly, a similar trend was also observed for metabolic activity **(Figure 4c)**. Moreover, the preservation of hMSC functionality in the presence of glycogen-loaded core-shell microgels compared to glycogen-free controls was clearly displayed by the continued secretion of glycogen degrading phosphorylase **(Figure 4d)** and pro-angiogenic VEGF **(Figure 4e)**. Together, these findings mirror the results obtained when glycogen was freely added to culture medium, indicating that entrapment of glycogen in microcapsules did not adversely affect its ability to release glucose nor its ability to sustain cell survival or function.

**Figure 4.**
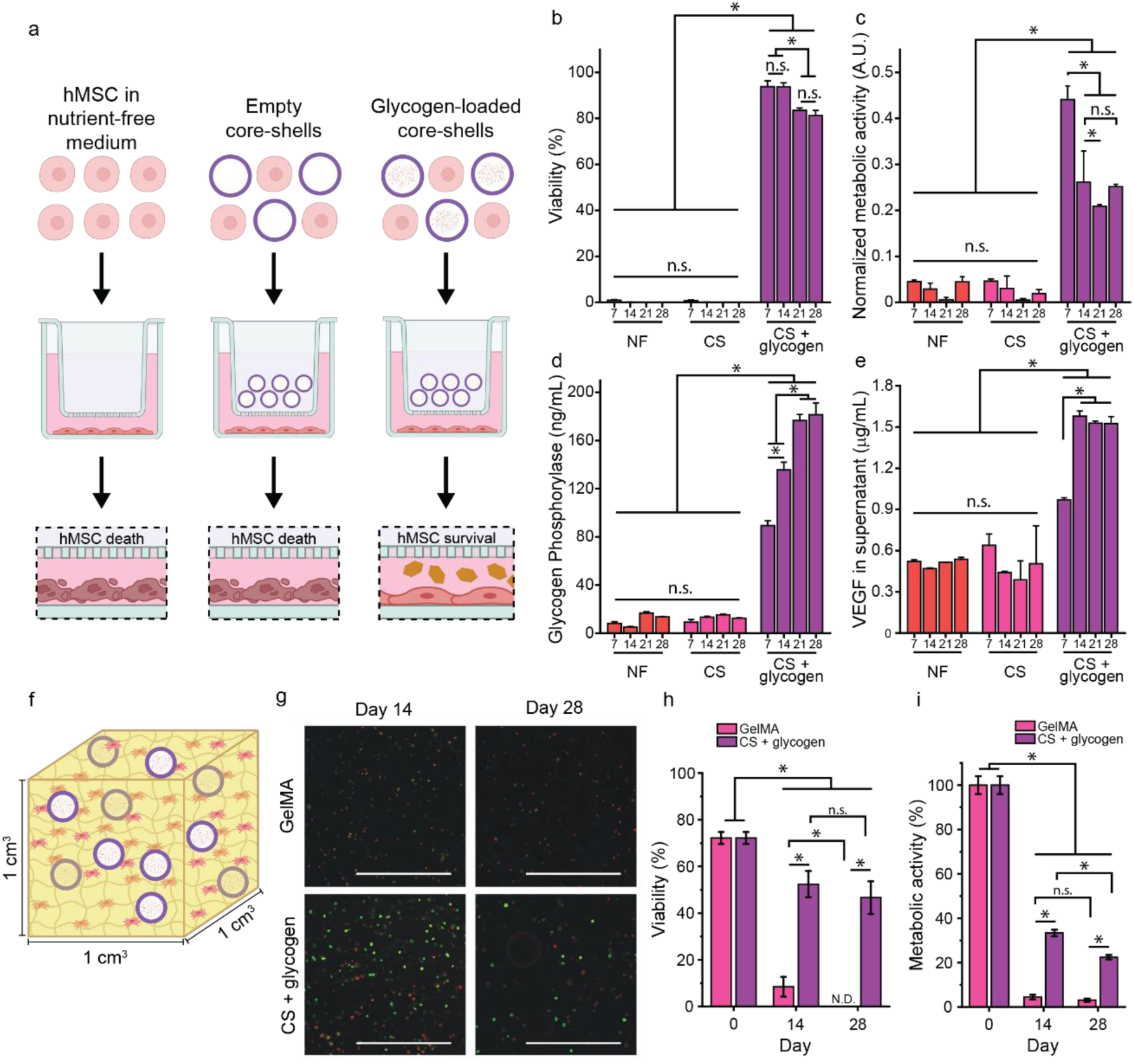
Glycogen-loaded core-shell microgels enabled self-feeding maintains hMSC function in large engineered constructs. a) schematic representation of transwell culture setup. b) Viability of hMSCs cultured in transwell system setup in anoxia with c) corresponding metabolic activity and d) glycogen phosphorylase (GP) and e) VEGF secretion. f) Schematic representation of 1 cm^3^ cell-laden and glycogen-loaded core-shell microgel containing constructs, g) representative live/dead images. scale bars: 1 mm h) with corresponding cell viability and i) metabolic activity measurements. n ≥ 3. Significance is indicated with * p < 0.05, and n.s. for p > 0.05, one-way ANOVA with Tukey post hoc test when normally distributed samples, otherwise Kruskal-Wallis.

Subsequently, the glycogen-containing microcapsules were incorporated into large, 1 cm^3^ sized hMSC-laden GelMA hydrogels to evaluate their potential to preserve the viability of large engineered tissue constructs **(Figure 4f)**. Again, the constructs were cultured in nutrient- and serum-free medium under anoxia to emulate an in vivo environment post-implantation, while cell-laden GelMA constructs served as controls. Viability drastically decreased for the empty core-shell microgel containing constructs after 14 days of anoxic incubation resulting in no detectable viable cells detected after 28 days of culture **(Figure 4g, h)**. Accordingly, the measured metabolic activity also drastically dropped after 14 days **(Figure 4i)**. In contrast, constructs containing glycogen-loaded core-shell microgels showed significantly higher cell viability over time compared to the control and remained stable after an initial decrease. Additionally, metabolic activity was preserved throughout the full 28-day culture period, further indicating functional maintenance of the engineered living materials. These results showcase that the cell-driven nutrient release from the homogeneously incorporated glycogen-based metabolic batteries overcomes the reliance of outside-in diffusion of systemic molecules such as nutrients or polysaccharide digesting enzymes (e.g. amylase). Consequently, we concluded that the glycogen-loaded core-shell microgels are a highly promising and versatile approach to engineer long-term self-feeding living materials using a nano-in-micro design strategy to effectively scale-up engineered living constructs towards cubic centimeter sizes.

### 2.5 Glycogen-based metabolic batteries accelerate integration of engineered constructs in vivo

To evaluate the potential of the glycogen-containing microcapsules as metabolic batteries in an in vivo setting, we investigated their potential to improve tissue regeneration and host-implant integration. To this end, glycogen containing microcapsules were incorporated into photocrosslinked GelMA hydrogels with or without 3×10^6^ hMSCs mL^-1^, which were subcutaneously implanted into nude mice for seven days **(Figure 5a)**. Additionally, constructs containing no additive, free glycogen, or glycogen-free microcapsules served as controls. To study the host-graft interactions, histological and immunohistochemical analyses were performed on midsagittal sections of constructs explanted one week post-implantation **(Figure 5b-e)**. Most importantly, hematoxylin and eosin (H&E) staining revealed that only the glycogen-containing microcapsules generated implants exhibiting an abundance of cells throughout the entire volume of the construct, while all other conditions only showed significant numbers of cells in their periphery **(Figure 5f)**. Interestingly, the cell-free nature of the implants revealed that the incorporation of microcapsules, regardless of glycogen presence, was sufficient to increase cellular infiltration into the implant. Improved cellular content also correlated with accelerated GelMA remodeling as well as de novo tissue formation in implants with glycogen-containing microcapsules **(Figure 5c)**. This was further corroborated by quantifying the remaining area of GelMA present within the slices, showing that implants with glycogen-containing microcapsules retained the least GelMA compared to all other conditions **(Figure 5g)**. In addition, glycogen-containing microcapsule implants were characterized by notably higher endothelial cell infiltration and neovascularization, as visualized using CD31+ staining, suggesting improved implant-host integration **(Figure 5d, h, i)**. In cell free-implants, supplementation of glycogen, both free and microcapsule entrapped, was sufficient to facilitate implant angiogenesis, while pristine cell-free implants remained completely void of blood vessel ingrowth. Surprisingly, supplementing implants with glycogen, both free and microencapsulated, significantly upregulated the number of CD206+ cells (i.e., pro-regenerative M2 macrophages), indicating the potential for metabolic materials to modulate immune responses in vivo **(Figure 5j).** Similar but more discrete effects were observed for hMSC-laden hydrogels **(Figure S4a–f)**, which were attributed to the small size of implanted constructs. increasing construct dimensions is expected to further amplify the benefits of the self-feeding strategy for living implants. Together, this suggests that endowing implants with self-feeding properties not only stimulates the survival and function of implanted cells, but also actively orchestrates host-implant interactions by stimulating host cells and thereby metabolically programming regenerative processes. Ultimately, glycogen-based metabolic batteries to enable self-feeding of engineered constructs may enable the fabrication of clinically relevant-sized engineered living materials, thereby offering a significant advancement in tissue engineering towards real-world, clinically relevant applications.

**Figure 5:**
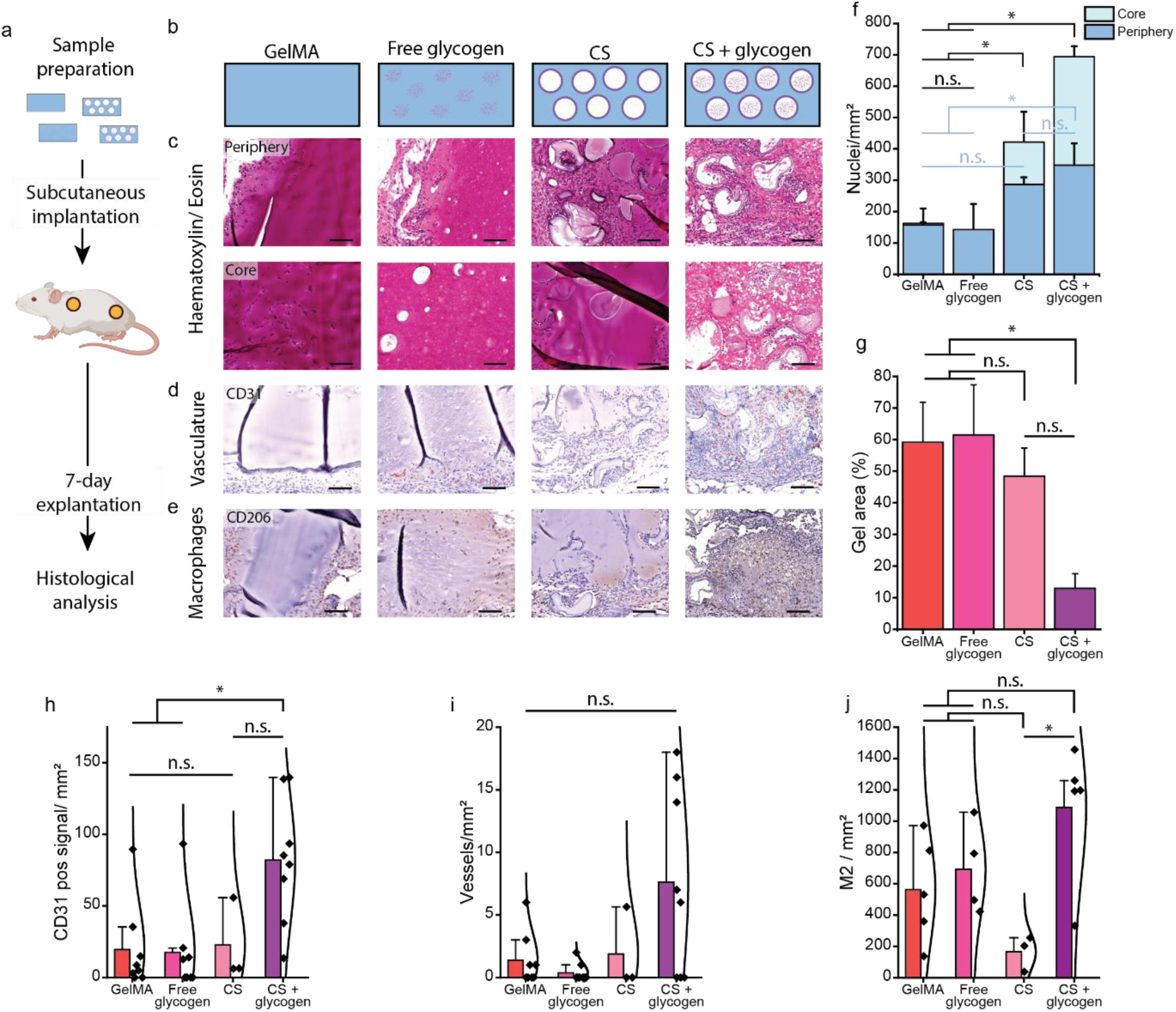
Glycogen-loaded core-shell microgels enabled self-feeding enhances engineered construct integration in vivo. a) schematic representation of subcutaneous implantation of the b) engineered constructs subcutaneously implanted into mice. c) Representative images of H&E stained midsagittal sections of the periphery and the core of hydrogels. d) representative images of CD31+ stained e) and CD206 stained sections. f) semi-quantification of the cellular infiltration, g) the remaining hydrogel area, h) the amount of CD31+ signal, i) the number of formed vessels, and j) M2 macrophages present. Scale bars: 200 µm. n ≥ 3. Significance is indicated with * p < 0.05, and n.s. for p > 0.05, one-way ANOVA with Tukey post hoc test when normally distributed samples, otherwise Kruskal-Wallis.

## 3. Conclusion

In this study, we introduce glycogen loaded core-shell microgels as a first-of-its-kind cell-responsive metabolic battery, leveraging nutritional nanoparticles that enable cell-mediated carbohydrate release to metabolically sustain engineered living materials. This innovative strategy offers prolonged nutrient release that preserves cell viability and function by supporting glycolysis for at least four weeks, even under hostile microenvironmental conditions such as anoxia, while simultaneously reducing the risk of implant acidosis. We demonstrate that human cells produce and secrete necessary enzymes to digest glycogen into metabolites that can be internalized and subsequently metabolized to support cellular energy demands. Moreover, we demonstrated that glycogen supplementation allows for sufficient energy generation to enable cells to secrete sufficient amounts glycogen-digesting enzymes, thereby establishing a cell-triggered and self-sustaining metabolic life-support system. Advantageously, this allows autonomous cell-centric nutrient release, wherein increases in cell number directly elevate nutrient availability, enabling engineered constructs to dynamically respond to the temporal needs of the engineered living matter. Additionally, the local secretion of glycogen degrading enzymes offers a novel avenue to locally produce nutrients without the reliance on outside-in diffusion of enzymes or nutrients. To make the use of glycogen universally applicable across diverse material systems, we developed a modular design strategy based on the production of glycogen-containing microcapsules. We demonstrated that these microcapsule effectively retained the nutritional nanoparticles for prolonged periods of time while allowing the free diffusion of glycogen-degrading enzymes and the resulting released glucose, thereby creating metabolic batteries that preserve cell viability and function for at least one month, even under anoxic conditions. The potential of this novel self-feeding strategy was evaluated for tissue engineering applications using subcutaneous implantations in mice. This revealed that engineered living materials containing glycogen-loaded microcapsules exhibited high cell viability, enhanced matrix remodeling, augmented tissue formation, pronounced M2 macrophage polarization, and accelerated vascularization. Interestingly, this demonstrates that incorporation of vast nutrient stores that can be controllably released has the potential to metabolically program regenerative processes within host tissues. Together, this establishes glycogen as an innovative method to maintain viability and tissue forming processes in large engineered living materials, with promising implications for numerous high-value applications including cultured meat, living soft robotics, drug screening models, and bioartificial organs.

## 4. Experimental Section/Methods

### List of materials and vendors

Minimal essential medium α with nucleosides, penicillin, streptomycin, trypsin-EDTA, GlutaMAX, d-glucose (dextrose) and Insulin-Transferrin-Selenium (ITS) 100X were purchased from Gibco. Basic fibroblast growth factor (bFGF) was purchased from Neuromics. Phosphate Buffered Saline (PBS) was purchased from Lonza. Dulbecco’s Modified Eagle’s Medium powder (DMEM powder without glucose, L-glutamine, phenol red, sodium pyruvate and sodium bicarbonate), glycogen from oyster, d-(-)-fructose, d-(+)-mannose, fluorescein isothiocyanate-dextran (average mol wt 2,000,000), Sodium Acetate, Sodium Butyrate, L-ascorbic acid 2-phosphate sesquimagnesium salt hydrate, palmitic acid, creatine monohydrate, L-aspartic acid, sodium pyruvate, α-ketoglutaric acid, linolenic acid, linoleic acid, oleic acid, amyloglucosidase from aspergillus niger, dopamine hydrochloride, sodium bicarbonate, Glyceraldehyde, n-hexadecane, Span80, hydrogen peroxide (H_2_O_2_), horseradish peroxidase (HRP), ascorbic acid (AsAP), calcein AM, ethidium homodimer-1 (EthD-1), fetal bovine serum (FBS), KB-228, Dulbecco’s Phosphate Buffered Saline (DPBS), Dextran, Glucose-6-Phosphate Assay Kit, Concanavalin A from Canavalia ensiformis, Lithium phenyl-2,4,6-trimethylbenzoylphosphinate, Millicell Cell Culture Inserts 24-well Hanging Inserts 3.0um PET, bovine serum albumin (BSA) Hematoxylin Gill #3 and Eosin-Y were purchased from Sigma-Aldrich/Merck. Dextran was conjugated with tyramine moieties as described by T. Kamperman et al. ^[50]^. Gelatin-Methacrylate (GelMA, 60% degree of functionalization, Df) was synthesized as described previously ^[62]^. Human phosphorylase glycogen liver form (PYGL) ELISA kit was purchased from Novus Biological. L-lactate assay kit was purchased from Abcam. Human VEGF-A (Vascular Endothelial Cell Growth Factor A) ELISA kit was purchased from Elabscience. PrestoBlue^TM^ cell viability reagent, 2″-[4-ethoxyphenyl]-5-[4-methyl-1- piperazinyl]-2,5″-bi-1H-benzimidazole) (NucBlue, Hoechst) and Nalgene Filtration Products were purchased from Thermo Fischer Scientific. Secondary antibodies against CD86 (antibodies.com), CD206 (Bio-Techne), CD31 (Cell Signaling Technology), and F4/80 (Bio-Rad) were purchased at various vendors. Fluorinated silane (Aquapel) was purchased from Vulcavite. Polydimethylsiloxane (PDMS, Sylgard 184) and Silicone Elastomer Curing Agent (Sylgard 184) were purchased from Dow Corning. Low pressure syringe pumps (neMESYS) were purchased from Cetoni. Gastight syringes (Hamilton) were purchased form IDEX Health & Science. Silicone tubing (inner diameter: 300μm, outer diameter: 640μm, thickness: 170μm) was purchased from Helix Medical.

### Cell isolation and expansion of hMSCs

Human mesenchymal stromal cells (hMSCs) were isolated from fresh bone marrow and cultured as previously described. In short, nucleated cells in the bone marrow aspirates were counted and subsequently seeded at 3000 cells cm^-2^ and cultured in culture medium consisting of 10% (v/v) FBS, 1% v/v, 100 U mL^−1^ Penicillin, 100 µg mL^−1^ Streptomycin, 1% (v/v) GlutaMax, 0.2 mᴍ ascorbic acid, and 1 ug L^-1^ bFGF (freshly added) in aMEM. The use of patient material was approved by the local ethical committee of the Medisch Spectrum Twente and informed written consent was obtained for all samples (METC\06003). At 80% confluency, hMSCs were detached using 0.25% trypsin-EDTA and either expanded, stored in liquid nitrogen, or used for experimentation.

### 2D cultures

For all 2D cultures, hMSCs were seeded at a density of 10000 cells cm^-2^ in culture medium and incubated overnight at 37 °C under 5% CO_2_. Subsequently, the culture medium was exchanged for medium composed of nutrient-free DMEM containing 3.7 g L^-1^ sodium bicarbonate, 1% v/v, 100 U mL^−1^ Penicillin, 100 µg mL^−1^ Streptomycin (denoted as Chemically Defined Medium) and the nutrient of interest. Afterwards, hMSCs were incubated in an oxygencontrolled environment (normoxia: 21% O_2_, and anoxia: 0.1% O_2_) at 37 °C under 5% CO_2_ until the end of the experiment. Depending on the experiment, d-glucose was added at various concentrations ranging from 0.06 – 10 g L^-1^. For the metabolic library, all nutrients were added directly to the medium at 5.5 mᴍ except for the fatty acids. Palmitic acid Linoleic acid, Linolenic acid and Oleic acid were dissolved at 3 mᴍ in 100% ethanol before adding it to the medium to reach a final concentration of 30 µᴍ.

### Cell viability and metabolic activity

Cell viability was assessed via incubation with 1.5 µᴍ calcein AM (live) and 6 µᴍ EthD-1 (dead) in DMEM and imaged using fluorescent microscopy. The presence of healthy, apoptotic and necrotic hMSCs was determined using the live/apoptotic/necrotic cells detection kit, following the manufacturer’s instructions. Metabolic activity was assessed by adding 10% PrestoBlue viability reagent directly to the medium, subsequent incubation and after which the fluorescent intensity of the supernatant was measured at using a plate read. Apoptotic, necrotic and healthy cells were imaged using the Apoptotic/Necrotic/Healthy cell detection kit according to the manufacturer’s instructions.

### Glycogen quantification

For the determination of glycogen remaining within the cell culture medium, cell culture supernatant was taken at 1, 3, 5, and 7 days, aliquoted and stored at -20°C until analyzed. For core-shell microgel containing experiments, 10 µL glycogen containing core-shell microgels were submerged in 500 µL PBS and incubated at 37°C for up to 14 days. At relevant timepoints (0, 1, 3, 7, 10, 14 days) 250uL aliquots were taken from the samples and stored at -20°C until analyzed, after which 250 µL fresh PBS was added again to the samples. Subsequently, the aliquots were mixed with amyloglucosidase (15 U mL^-1^) in MilliQ water at a 1:1 (v/v) ratio and incubated for 30 min at 37°C. Subsequently, the aliquot-enzyme mixture was combined 1:1 with glucose oxidase (400 U mL^-1^) and incubated at room temperature for 20 min. Finally, 10 μL of the resulting sample solution was mixed with 40 µL of 50 mᴍ sodium phosphate in PBS at pH 7.4, and 50 μL Ampliflu Red-HRP solution, resulting in a final concentration of 0.2 U mL^-1^ HRP and 100 μᴍ Ampliflu Red. This mixture was further incubated for 30 min at room temperature, protected from light, after which fluorescence intensity was measured using a VarioskanLUX plate reader (Thermo Fischer Scientific) at excitation/emission wavelengths of 530/590 nm.

### Glucose quantification

A d-glucose standard was prepared in PBS ranging from 0 – 1 g L^-1^. Standard and samples were mixed 1:1 with a reaction buffer composed of 20 U mL^-1^ HRP, 200 U mL^-1^ GOX and 0.5x DAB in PBS and incubated for 20 minutes at room temperature, protected from light. Subsequently, optical density was measured at 504 nm in a plate reader (Multiscan GO, Thermo Fisher Scientific).

### Lactate quantification

hMSCs were cultured under anoxia in nutrient- and serum-free medium either containing 1 g L^-1^ glycogen or glucose serving as control. After 1, 3, 5, and 7 days the supernatant was collected and stored at -20°C until analyzed. L-lactate was then quantified based on the manufacturer’s instructions.

### Glycogen phosphorylase quantification

hMSCs (5k, 10k, or 20k cells per well) were cultured under anoxia in chemically defined medium containing glycogen (0, 1, and 10 g L^-1^) for a maximum of 14 days. Supernatant was collected at various timepoints and stored at -20°C. The glycogen phosphorylase content was quantified using an ELISA kit following manufacturer’s protocol (NovusBio).

### Glucose-6-phosphate quantification

Glycogen solutions (0, 1, and 10 g L^-1^) and glycogen-loaded core-shell microgels (0, 30, 50, and 100 g L^-1^) were exposed to conditioned medium containing glycogen-phosphorylase for glycogen breakdown into glucose-1-phosphate. Subsequently, the samples were exposed to 5 U mL^-1^ of phoshoglucomutase to induce glucose-6-phosphate (G6P) conversion. The concentration of G6P was quantified using a G6P Assay following manufacturer’s protocol (Merck)

### Dex-TA synthesis

Dextran (MW 40 kg mol^−1^) was functionalized with tyramine as previously described.^[63]^ Dextran-tyramine (Dex-TA) contained ≈14 tyramine moieties per 100 repetitive monosaccharide units.

### Osmolarity

The osmolarity of glycogen and glucose samples in PBS or medium was measured using a Semi-Micro Osmometer (K-7400S, Knauer).

### Glycogen particle size

Dynamic light scattering size measurements were performed with the Malvern Zetasizer. A solution of 0.025 g L^-1^ glycogen in ddH_2_O was pipetted into a DLS cuvette which was placed in the Zetasizer. For both size and charge measurements, the following settings were used: optical absorbance 0.2, peak at 450nm and a refractive index of 1.687. Each sample was measured 5 times with 15 runs per measurement.

Additionally, size was analyzed using transmission electron microscopy. Briefly, glycogen was deposited on a carbon-coated copper grid and negatively stained using 2% uranyloxalate-acetate (pH 7). Glycogen nanoparticles were then imaged using a transmission electron microscope (FEI Tecnai 12, Thermo Fisher Scientific) and the size of 20 single particles were analyzed using ImageJ. *Blocking glycogen phosphorylase:* hMSCs were seeded in 24 wells plates as previously described in nutrient- and serum-free medium containing 0, 0.1, and 1 g L^-1^ glycogen and 0 or 100 µᴍ of glycogen phosphorylase blocker (KB228). 0 or 100 µᴍ final concentration of fresh blocker was added daily. Cell viability was assessed after three days under anoxia as previously described.

### VEGF secretion and gene expression

VEGF secretion was measured by analyzing the hMSC supernatant with a commercial VEGF-A ELISA kit, following the manufacturer’s instructions. For the gene expression, hMSCs were cultured as previously described in nutrient- and serum-free medium containing 0, 0.01, 0.1, 1 g L^-1^ glycogen and cultured for 7 days under anoxia. Total RNA was extracted by all cells using TRIzol and immediately storing the lysate at -80°C until further use. RNA was then isolated using the miRNeasy kit following the manufacturer’s instructions. Subsequently, cDNA was synthesized using the iScript cDNA synthesis kit. cDNA was then subjected to qPCR using SensiMix SYBR & fluorescein kit on a CFX Connect Real-time System (Bio-Rad). Gene expression was normalized on beta-actin (ACTB) expression. ACTB was validated to act as a stable reference gene within the experiment’s dataset. Three biological replicates were carried out per time-point.

### Fluorescent labelling of glycogen (glycogen-AF488)

A fluorescent dye (5-(4,6-dichlorotriazinyl)aminofluorescein (5-DTAF)) was covalently labelled to glycogen. Specifically, 250 mg of oyster glycogen was first dissolved in 25 mL of MilliQ water and 0.1 ᴍ Na_2_CO_3_ solution was used to tune the pH to 9.5. Then 380 µL of 5-DTAF (1 mg mL^-1^ stock in anhydrous DMSO, 0.125 mg oyster glycogen, to label 0.05 % of repeating glucose units) was added in a dropwise manner while maintaining vigorous stirring. The reaction was proceeded at 25 °C in a dark environment for four hours. The solution was then precipitated in ice cold ethanol and subsequently washed three times, followed by drying under vacuum. The recovered solid was redissolved in MilliQ water, and dialyzed (MWCO 3500) for three days. Finally, the dialyzed solution was freeze-dried and stored for subsequent use.

### Core-shell preparation

The mold for the microfluidic droplet generator was printed with stereolithography (Formlabs 3B+ SLA 3D printer) using Clear V4 resin (Formlabs). The microfluidic droplet generator was subsequently produced using PDMS, and plasma treated for proper bonding to microscope slides. Prior to core-shell microgel production, the microfluidic droplet generator was perfused with Aquapel to ensure hydrophobicity of all components. Microdroplets were created by emulsifying a hydrogel precursor solution containing 10% w/v tyramine conjugated dextran (Dex-TA), glycogen (0 – 100 g L^-1^), and 80 U mL^-1^ horseradish peroxidase (HRP) in N-hexadecane containing 1% (v/v) Span-80 surfactant at a 10:80 µL min^-1^ (hydrogel precursor:oil) flow ratio. Flowing the microdroplets through a semipermeable silicone tubing that was submerged in a hydrogen peroxide bath (30% w/w) allowed for an outside-in crosslink of the microdroplets. After washing the solution with pure N-hexadecane followed by breaking the emulsion with sterile PBS, this resulted in the formation of a crosslinked hydrogel shell surrounding non-crosslinked hydrogel precursor solution. The core-shell microgels were stained using EthD-1 (4 x 10^-6^ ᴍ) to stain the Dex-TA and fluorescently labelled glycogen was mixed 1:10 with unlabeled glycogen to visualize the glycogen.

Stained core-shell microgels were analyzed using fluorescence microscopy (EVOS FL), confocal fluorescence microscopy (Zeiss LSM 510), and fluorescence recovery after photobleaching (FRAP; Nikon A1+). The FRAP curve was generated by plotting the background-subtracted fluorescence of the bleached region over time, normalized to the bleach-rate–corrected mean intensity measured before bleaching. The bleach rate was calculated from the fluorescence in an unbleached area of the sample (outside the bleach spot), normalized to its pre-bleach mean intensity, as previously described.^[63]^

### FITC-labeled dextran release from core-shell microgels

FITC-labeled dextran (Dex-FITC; 10, 70, 300, and 500 kDa) and fluorescently labelled glycogen were encapsulated within the core-shell microgels during production. Release from the core-shell microgels, quantified as the loss of fluorescence over time, was measured in real-time by time-lapse fluorescence imaging (EVOS FL Auto 2).

### Transwell experiments

hMSCs were seeded in 24 wells plates as previously described. Instead of adding nutrients directly to the nutrient- and serum-free medium, 10 µL of core-shell microgels were added to a transwell insert that was added to the respective wells. Cell viability and metabolic activity were assessed as previously described.

### 3D tissue constructs

Large, 1 cm^3^, constructs were created by mixing 3 x 10^6^ mL^-1^ hMSCs with 7.5 wt% GelMA, 0.1 wt% LAP, and, when stated, 25% (i.e. 250 µL) of glycogen-loaded core-shell microgels. The solution was transferred to an in-house manufactured 1 cm^3^ mold, and crosslinked using UV light for two minutes (JD818 UV-lamp, Jiadi). The constructs were incubated overnight in complete aMEM culture medium to allow the cells to recover, after which they were placed in anoxia with chemically-defined medium with 10% FBS. Cell viability and metabolic activity were assessed on day 14 and day 28 as previously described.

### In vivo implantation

To evaluate the in vivo behavior of self-feeding tissues, 5% (w/v) GelMA hydrogels containing free glycogen (10 g L^-1^), empty core-shell microgels and glycogen loaded core-shell microgels were fabricated with and without hMSCs (3 × 10^6^ cells mL^-1^) hMSCs. Hemispherical hydrogels (1 mm thickness and 8 mm diameter) were subcutaneously implanted in the backs on 12-week-old nude rats (NIH-Foxn1^rnu^ Rat, Charles River). Animal procedures of subcutaneous test (e.g., hydrogel implantation in nude rats) were approved by the Institutional Animal Use and Care Committee of Harvard Medical School (Protocol number: 2017N000114). Briefly, anesthesia was induced and maintained with isoflurane in spontaneously breathing rats. After explantation, samples were fixed in formalin and embedded in paraffin.

### Histological analysis

The paraffin embedded samples were cut into 5 um thin slices using a microtome. These tissue sections were then stained for histological analysis. For the analysis of the integration of the implanted tissues, the slices were rehydrated and stained with hematoxylin (Gill #3; Sigma-GHS332) for 10 min, and Eosin-Y (Sigma-HT110132) for 30 s.

For the immunostainings, antigen retrieval was performed by boiling the slides for 12 minutes in citrate buffer. Slides for immunohistochemical staining were blocked with 0.3% H_2_O_2_ for 10 min, and 5 w/v % PBS-BSA in PBS at room temperature for 1 hour. CD31 (1:100) and CD206 (3.3 ug mL^-1^) were incubated overnight at 4°C. After washing, goat-anti-mouse secondary antibody conjugated to HRP (Agilent P0447) was added and incubated at room temperature for 1 hour, and the staining was visualized using DAB (Abcam ab64238). Stained histological sections were imaged (Hamamatsu Nanozoomer). The semi-quantitative analysis for the remaining GelMA area, CD31 positive area and the number of macrophages was performed on at least three areas per slide and performed using Fiji ImageJ.

### Statistical analysis

Statistics were performed using OriginPro software. Data were represented as mean ± standard deviation and statistical significance was determined at p<0.05 (*) using one-way ANOVA with Tukey post hoc test for normally distributed datasets, otherwise Kruskal-Wallis was used. Schematics were created using bioRender.com and Adobe Illustrator.

## Acknowledgements

J.L acknowledges financial support from the European Research Council (Starting Grant, 759425). The authors thank Sara Gil Izquierdo for excellent technical assistance with the TEM imaging.

## References

[1] Elkhoury, K., Sanchez-Gonzalez, L., Lavrador, P., et al., “Gelatin methacryloyl (GelMA) nanocomposite hydrogels embedding bioactive naringin liposomes,” Polymers 12, no. 12 (2020).

[2] Zagury, Y., Ianovici, I., Landau, S., Lavon, N., Levenberg, S., “Engineered marble-like bovine fat tissue for cultured meat,” Communications Biology 5, no. 1 (2022).

[3] Farzin, A., Hassan, S., Moreira Teixeira, L. S., et al., “Self-Oxygenation of Tissues Orchestrates Full-Thickness Vascularization of Living Implants,” Advanced functional materials 31, no. 42 (2021).

[4] Appiah, C., Arndt, C., Siemsen, K., Heitmann, A., Staubitz, A., Selhuber-Unkel, C., “Living Materials Herald a New Era in Soft Robotics,” Advanced Materials 31, no. 36 (2019).

[5] Owens, R. M., “Advanced tissue engineering for in vitro drug safety testing,” MRS Communications 13, no. 5 (2023).

[6] Rouwkema, J., Rivron, N. C., van Blitterswijk, C. A., “Vascularization in tissue engineering,” Trends Biotechnol 26, no. 8 (2008).

[7] McMurtrey, R. J., “Analytic models of oxygen and nutrient diffusion, metabolism dynamics, and architecture optimization in three-dimensional tissue constructs with applications and insights in cerebral organoids,” Tissue Engineering Part C: Methods 22, no. 3 (2016).

[8] Rouwkema, J., Koopman, B. F., Blitterswijk, C. A. V., Dhert, W. J., Malda, J., “Supply of nutrients to cells in engineered tissues,” Biotechnology and Genetic Engineering Reviews 26, no. 1 (2009).

[9] Rustad, K. C., Sorkin M Fau - Levi, B., Levi B Fau - Longaker, M. T., Longaker Mt Fau - Gurtner, G. C., Gurtner, G. C., “Strategies for organ level tissue engineering,” no. 1555-8592 (Electronic) (2010).

[10] Pedraza, E., Coronel, M. M., Fraker, C. A., Ricordi, C., Stabler, C. L., “Preventing hypoxia-induced cell death in beta cells and islets via hydrolytically activated, oxygen-generating biomaterials,” Proceedings of the National Academy of Sciences 109, no. 11 (2012).

[11] Willemen, N. G., Hassan, S., Gurian, M., et al., “Enzyme-mediated alleviation of peroxide toxicity in self-oxygenating biomaterials,” Advanced healthcare materials 11, no. 13 (2022).

[12] Erdem, A., Darabi, M. A., Nasiri, R., et al., “3D bioprinting of oxygenated cell-laden gelatin methacryloyl constructs,” Advanced healthcare materials 9, no. 15 (2020).

[13] Rizzo, S. A., Bartley, O., Rosser, A. E., Newland, B., “Oxygen-glucose deprivation in neurons: implications for cell transplantation therapies,” Progress in Neurobiology 205(2021).

[14] Deschepper, M., Oudina, K., David, B., et al., “Survival and function of mesenchymal stem cells (MSCs) depend on glucose to overcome exposure to long-term, severe and continuous hypoxia,” Journal of cellular and molecular medicine 15, no. 7 (2011).

[15] Deschepper, M., Manassero, M., Oudina, K., et al., “Proangiogenic and prosurvival functions of glucose in human mesenchymal stem cells upon transplantation,” Stem Cells 31, no. 3 (2013).

[16] Moya, A., Paquet, J., Deschepper, M., et al., “Human mesenchymal stem cell failure to adapt to glucose shortage and rapidly use intracellular energy reserves through glycolysis explains poor cell survival after implantation,” Stem Cells 36, no. 3 (2018).

[17] Coyle, R., Yao, J., Richards, D., Mei, Y., “The effects of metabolic substrate availability on human adipose-derived stem cell spheroid survival,” Tissue Engineering Part A 25, no. 7-8 (2019).

[18] Fois, M. G., Zengin, A., Song, K., et al., “Nanofunctionalized Microparticles for Glucose Delivery in Three-Dimensional Cell Assemblies,” ACS Applied Materials & Interfaces 16, no. 14 (2024).

[19] Zargarzadeh, M., Silva, A. S., Nunes, C., Coimbra, M. A., Custódio, C. A., Mano, J. F., “Self-glucose feeding hydrogels by enzyme empowered degradation for 3D cell culture,” Materials Horizons 9, no. 2 (2022).

[20] Lescher, A., Kansou, K., Della Valle, G., Petite, H., Lourdin, D., “Evaluation of extruded starch foam for glucose-supplying biomaterials,” Carbohydrate Polymers (2024).

[21] Fastman, N. M., Liu, Y., Ramanan, V., et al., “The structural mechanism of human glycogen synthesis by the GYS1-GYG1 complex,” Cell reports 40, no. 1 (2022).

[22] Adeva-Andany, M. M., González-Lucán, M., Donapetry-García, C., Fernández-Fernández, C., Ameneiros-Rodríguez, E., “Glycogen metabolism in humans,” BBA clinical 5(2016).

[23] Kanungo, S., Wells, K., Tribett, T., El-Gharbawy, A., “Glycogen metabolism and glycogen storage disorders,” Annals of translational medicine 6, no. 24 (2018).

[24] Besford, Q. A., Cavalieri, F., Caruso, F., “Glycogen as a building block for advanced biological materials,” Advanced materials 32, no. 18 (2020).

[25] Gopinath, V., Saravanan, S., Al-Maleki, A., Ramesh, M., Vadivelu, J., “A review of natural polysaccharides for drug delivery applications: Special focus on cellulose, starch and glycogen,” Biomedicine & Pharmacotherapy 107(2018).

[26] Gálisová, A., Jirátová, M., Rabyk, M., et al., “Glycogen as an advantageous polymer carrier in cancer theranostics: Straightforward in vivo evidence,” Scientific Reports 10, no. 1 (2020).

[27] Chandel, N. S., “Glycolysis,” Cold Spring Harbor Perspectives in Biology 13, no. 5 (2021).

[28] Emanuelle, S., Brewer, M. K., Meekins, D. A., Gentry, M. S., “Unique carbohydrate binding platforms employed by the glucan phosphatases,” Cellular and molecular life sciences 73(2016).

[29] Sharma, V., Ichikawa, M., Freeze, H. H., “Mannose metabolism: more than meets the eye,” Biochemical and biophysical research communications 453, no. 2 (2014).

[30] Sun, S. Z., Empie, M. W., “Fructose metabolism in humans–what isotopic tracer studies tell us,” Nutrition & metabolism 9(2012).

[31] Legrand, P., Rioux, V., “The complex and important cellular and metabolic functions of saturated fatty acids,” Lipids 45(2010).

[32] Gregory, M. K., King, H. W., Bain, P. A., Gibson, R. A., Tocher, D. R., Schuller, K. A., “Development of a fish cell culture model to investigate the impact of fish oil replacement on lipid peroxidation,” Lipids 46(2011).

[33] Schönfeld, P., Wojtczak, L., “Short-and medium-chain fatty acids in energy metabolism: the cellular perspective,” Journal of lipid research 57, no. 6 (2016).

[34] Stegen, S., Van Gastel, N., Eelen, G., et al., “HIF-1α promotes glutamine-mediated redox homeostasis and glycogen-dependent bioenergetics to support postimplantation bone cell survival,” Cell metabolism 23, no. 2 (2016).

[35] Eales, K. L., Hollinshead, K. E., Tennant, D. A., “Hypoxia and metabolic adaptation of cancer cells,” Oncogenesis 5, no. 1 (2016).

[36] Chinopoulos, C., “Which way does the citric acid cycle turn during hypoxia? The critical role of α-ketoglutarate dehydrogenase complex,” Journal of neuroscience research 91, no. 8 (2013).

[37] Smirnoff, N., “Ascorbic acid metabolism and functions: A comparison of plants and mammals,” Free Radical Biology and Medicine 122(2018).

[38] Santacruz, L., Arciniegas, A. J. L., Darrabie, M., et al., “Hypoxia decreases creatine uptake in cardiomyocytes, while creatine supplementation enhances HIF activation,” Physiological reports 5, no. 16 (2017).

[39] Engl, E., Garvert, M. M., “A prophylactic role for creatine in hypoxia?,” Journal of Neuroscience 35, no. 25 (2015).

[40] Güemes, M., Rahman, S. A., Hussain, K., “What is a normal blood glucose?,” Archives of disease in childhood 101, no. 6 (2016).

[41] Listenberger, L. L., Ory, D. S., Schaffer, J. E., “Palmitate-induced apoptosis can occur through a ceramide-independent pathway,” Journal of Biological Chemistry 276, no. 18 (2001).

[42] Jane B. Reece, L. A. U., Michael L. Cain, Steven A. Wasserman, Peter V. Minorsky, Robert B. Jackson 2010, Cellular Respiration and Fermentation. ISBN 0321558235.

[43] Date, K., Satoh, A., Iida, K., Ogawa, H., “Pancreatic alpha-Amylase Controls Glucose Assimilation by Duodenal Retrieval through N-Glycan-specific Binding, Endocytosis, and Degradation,” J Biol Chem 290, no. 28 (2015).

[44] Tiffon, C., “Defining Parallels between the Salivary Glands and Pancreas to Better Understand Pancreatic Carcinogenesis,” Biomedicines 8, no. 6 (2020).

[45] Davydiuk, N., Londhe, V., Maitz, M. F., Werner, C., Fery, A., Besford, Q. A., “The interaction of glycogen nanoparticles with human blood,” Nanoscale 17, no. 1 (2025).

[46] Fernandes, S., Xu, R., De Rose, R., et al., “Trafficking Glycogen Nanoparticles through Lymph Node Tissues for the Delivery of Small and Large Bioactive Molecules,” ACS Nano 19, no. 44 (2025).

[47] Lavogina, D., Lust, H., Tahk, M.-J., et al., “Revisiting the resazurin-based sensing of cellular viability: Widening the application horizon,” Biosensors 12, no. 4 (2022).

[48] Hopp, A.-K., Grüter, P., Hottiger, M. O., “Regulation of glucose metabolism by NAD+ and ADP-ribosylation,” Cells 8, no. 8 (2019).

[49] Perlman, J. M., Volpe, J. J. 2018, “Chapter 25 - Glucose.” in Volpe’s Neurology of the Newborn (Sixth Edition). edited by Volpe, J. J., Inder, T. E., Darras, B. T., et al.: Elsevier. ISBN 978-0-323-42876-7.

[50] Paredes-Flores, M. A., Rahimi, N., Mohiuddin, S. S. 2024, “Biochemistry, glycogenolysis.” in StatPearls [Internet]. StatPearls Publishing.

[51] Young, F. G., “Claude Bernard and the discovery of glycogen,” British Medical Journal 1, no. 5033 (1957).

[52] Liu, Q.-H., Tang, J.-W., Wen, P.-B., Wang, M.-M., Zhang, X., Wang, L., “From prokaryotes to eukaryotes: insights into the molecular structure of glycogen particles,” Frontiers in molecular biosciences 8(2021).

[53] Bras, N. F., Fernandes, P. A., Ramos, M. J., “Understanding the Rate-Limiting Step of Glycogenolysis by Using QM/MM Calculations on Human Glycogen Phosphorylase,” ChemMedChem 13, no. 15 (2018).

[54] Park, K.-Y., Ay, I., Avery, R., et al., “New biomarker for acute ischaemic stroke: plasma glycogen phosphorylase isoenzyme BB,” *Journal of Neurology*, Neurosurgery & Psychiatry 89, no. 4 (2018).

[55] Ghimire, A., Giri, S., Khanal, N., et al., “Diagnostic accuracy of glycogen phosphorylase BB for myocardial infarction: A systematic review and meta-analysis,” Journal of Clinical Laboratory Analysis 36, no. 5 (2022).

[56] Krock, B. L., Skuli, N., Simon, M. C., “Hypoxia-induced angiogenesis: good and evil,” Genes & cancer 2, no. 12 (2011).

[57] Bhowmick, T., Biswas, S., Mukherjee, A., “Cellular response during cellular starvation: A battle for cellular survivability,” Cell Biochemistry and Function 42, no. 5 (2024).

[58] Zois, C., Harris, A., “Glycogen metabolism has a key role in the cancer microenvironment and provides new targets for cancer therapy,“ Journal of molecular medicine (Berlin, Germany) 94(2016).

[59] Van Loo, B., Salehi, S., Henke, S., et al., “Enzymatic outside-in cross-linking enables single-step microcapsule production for high-throughput three-dimensional cell microaggregate formation,” Materials Today Bio 6(2020).

[60] Besford, Q. A., Weiss, A. C. G., Schubert, J., et al., “Protein Component of Oyster Glycogen Nanoparticles: An Anchor Point for Functionalization,” ACS Appl Mater Interfaces 12, no. 35 (2020).

[61] Artigas, A. C., Barón, C., Parody-Morreale, A., “Molecular studies of glycogen phosphorylase b from bovine liver,” Int J Biol Macromol 17, no. 2 (1995).

[62] Zhang, Y. S., Davoudi, F., Walch, P., et al., “Bioprinted thrombosis-on-a-chip,” Lab on a Chip 16, no. 21 (2016).

[63] Kamperman, T., Willemen, N. G., Kelder, C., et al., “Steering Stem Cell Fate within 3D Living Composite Tissues Using Stimuli-Responsive Cell-Adhesive Micromaterials,” Advanced Science 10, no. 10 (2023).

