## Supplemental File for "Self-feeding living materials enabled by cell responsive glycogen nanoparticles as metabolic batteries"

*Glycogen uptake:* hMSCs and RAW cells were seeded into ibidi slides and incubated over night under standard culture conditions (21% $O_2$, at 37 °C under 5% $CO_2$) for attachment. Afterwards, cells were washed in PBS and chemically defined medium with and without glycogen-AF488 (1 g $L^{-1}$) was added. After 24 hours of incubation at 37 °C under 5% $CO_2$, cells were fixed in formalin, stained with phalloidin and DAPI and imaged using confocal microscopy.
*Glycogen Stability:* 2 g $L^{-1}$ oyster glycogen was dissolved in MilliQ and incubated at 37 °C for 28 days. After 0, 14, and 28 days, the molecular weight (Mw) was measured using static light scattering. Toluene was used as standard and glycogen concentrations of 0.1, 0.5, 1, 1.5 and 2 g $L^{-1}$ were used to obtain a Debye plot from which the Mw was derived.
*GP secretion of different cell types:* Various cell types were expanded towards 80% confluency after which supernatant was collected. Afterwards, GP was quantified using an ELISA kit following manufacturer's protocol (Elabscience).
*Reperfusion differentiation:* hMSCs were cultured in 2D as described in the main manuscript under anoxia in chemically defined medium containing only 1 g $L^{-1}$ of glycogen. After 28 days of continuous cultured without medium refresh, differentiation medium was directly added to the cells and they were placed moved into normoxia (21% $O_2$, at 37 °C under 5% $CO_2$). After 24 hours, medium was refreshed and differentiation continued for 21 days with 3 medium changes per week. For osteogenic differentiation, the medium consisted of aMEM containing 10% (v/v) FBS, 1% v/v, 100 U $mL^{-1}$ Penicillin, 100 µg $mL^{-1}$ Streptomycin, 1% (v/v) GlutaMax, 0.2 mM ascorbic acid, 10 mM beta-glycerophosphate (freshly added) and 10 nM dexamethasone (freshly added). Osteogenic differentiation was then visualized using brightfield microscopy after staining the samples with 20 g L−1 Alizarin Red S in saline dH2O.
For adipogenic differentiation, the medium consisted of DMEM containing 10% (v/v) FBS, 1% v/v, 100 U $mL^{-1}$ Penicillin, 100 µg $mL^{-1}$ Streptomycin, 1% (v/v) GlutaMax. Each medium refresh 1 µM dexamethasone, 0.5 mM IBMX, 0.2 mM Indomethacin and 10 µg $mL^{-1}$ insulin were freshly added. Adipogenic differentiation was visualized using brightfield microscopy after staining the samples with 1.8 g L−1 Oil Red O in a 2-propanol/PBS mixture (3:2).

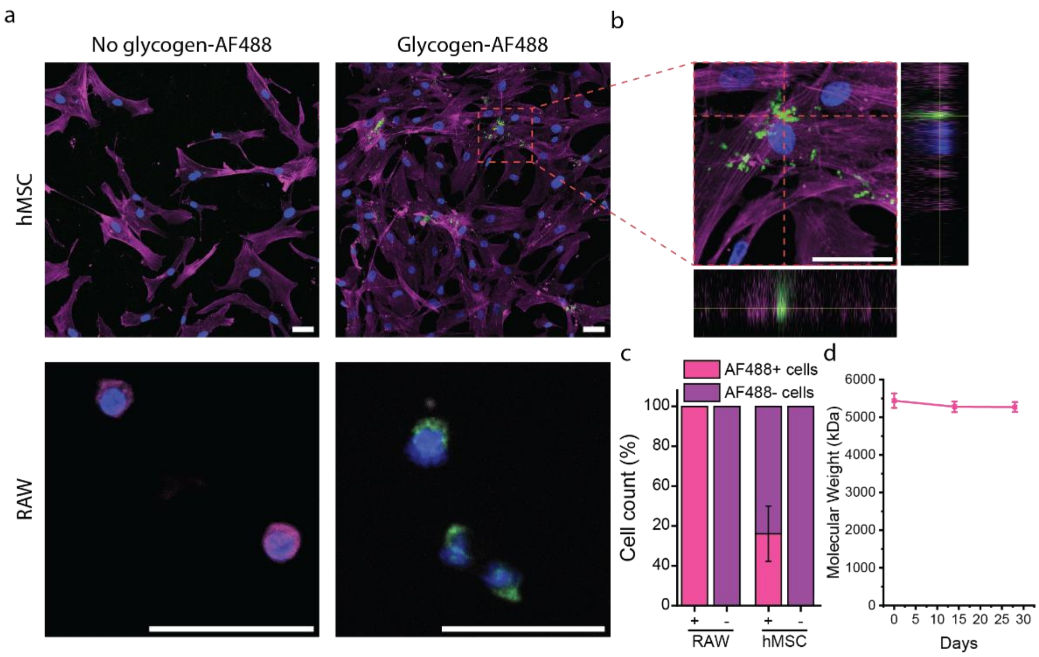


**Figure S1:** a) Representative images of hMSCs and RAW cells incubated without and with glycogen-AF488. b) Height-resolved confocal fluorescent micrograph of hMSCs showcasing the internalization of glycogen-AF488. c) Quantification of the fraction of cells that internalized glycogen-AF488. Scale bars: 50 µm. d) Molecular weight of glycogen incubated in PBS at 37 °C for a period of 30 days.

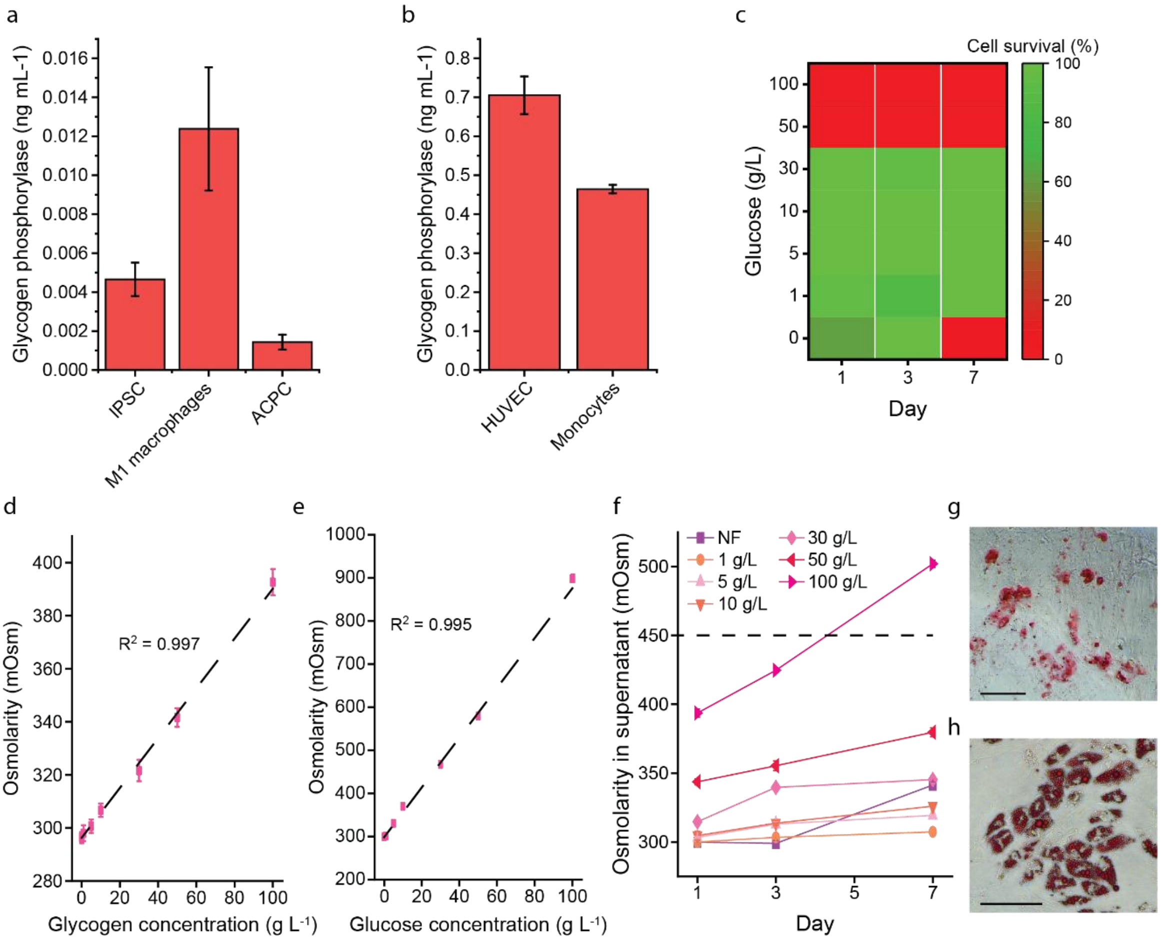


**Figure S2.** a) Glycogen phosphorylase secretion of induced pluripotent stem cells (IPSC), M1 polarized macrophages, and articular cartilage progenitor cells (ACPC), b) as well as human umbilical cord endothelial cells (HUVEC) and CD14+ monocytes. c) hMSC viability after exposure to various glucose concentrations d) Effect of glycogen and e) glucose supplementation to PBS. f) Osmolarity of culture medium supplemented with various initial glycogen concentrations over time. g) Alizarin red stained and h) oil red O stained hMSCs differentiated for three weeks towards either an osteogenic or adipogenic lineage after being cultured in in nutrient- and serum-free medium supplemented with glycogen under anoxic conditions for 28 days. Scale bars: 100 µm.

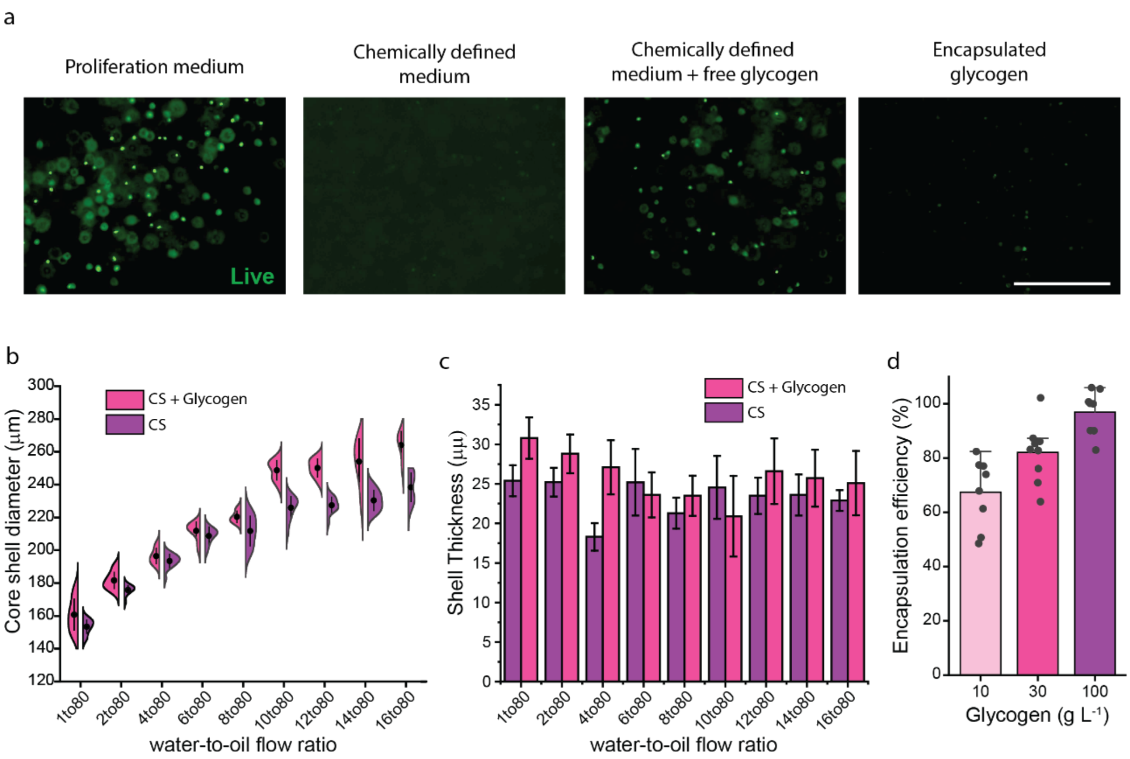


**Figure S3.** a) Representative images of viable hMSCs in Dex-TA with various nutrient supplements, scale bar: 400µm b) Diameter of glycogen loaded and empty core-shell microgels at different water to oil flow ratios and c) their corresponding shell thickness. d) Encapsulation efficiency of glycogen within core-shell microgels.

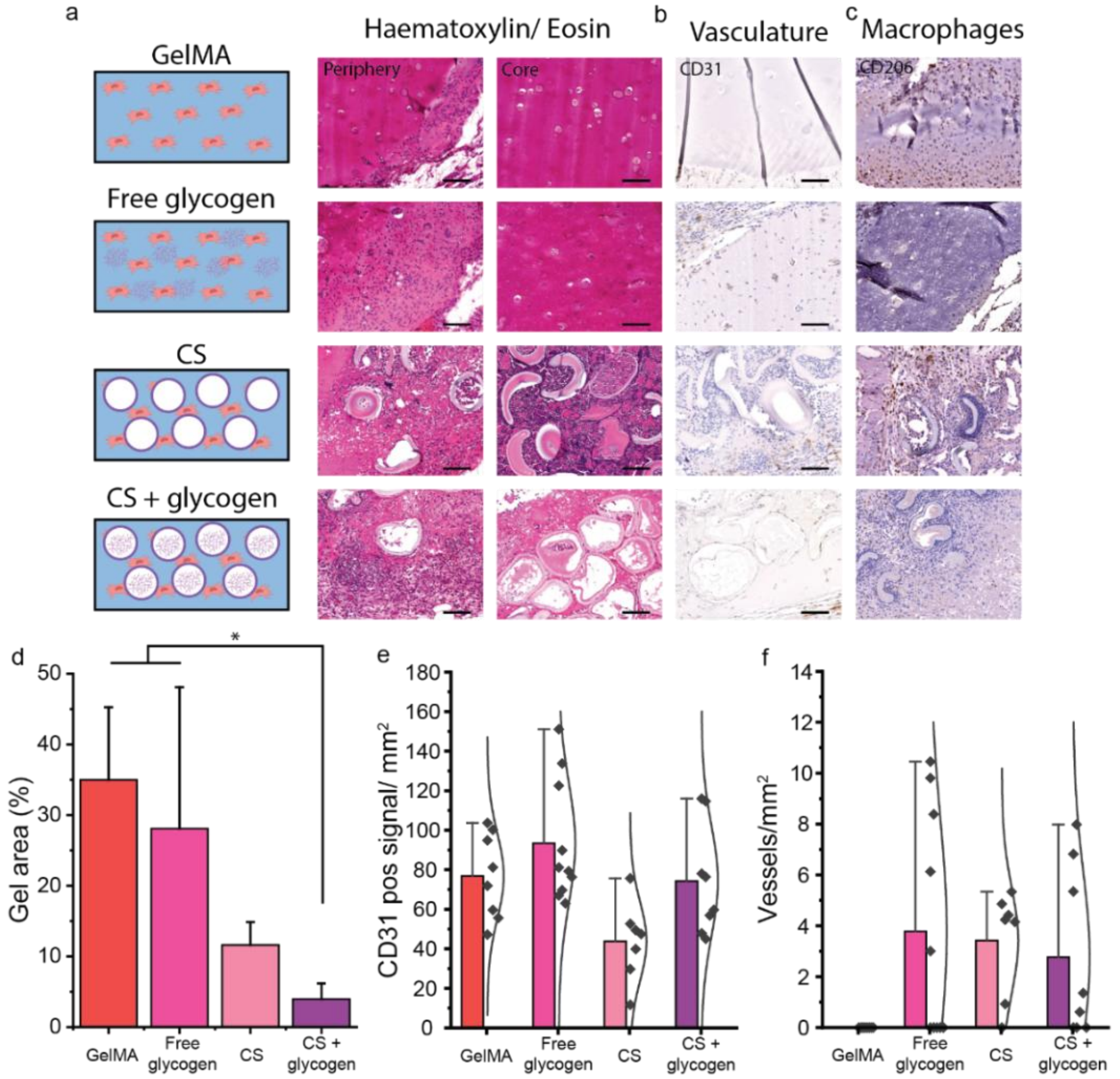


.

**Figure S4.** a) Representative images of H&E stained midsagittal sections of the periphery and the core of hMSC-laden hydrogels. b) representative images of CD31+ stained c) and CD206 stained sections. d) the amount of CD31+ signal, e) the number of formed vessels, f) M2 macrophages present. Scale bars: 200 µm; $n \geq 3$ Significance is indicated with * $p < 0.05$, and n.s for $p > 0.05$, one-way ANOVA with Tukey post hoc test when normally distributed samples, otherwise Kruskal-Wallis.
